# Platelet Membrane-Coated SPION for Targeted Paclitaxel Delivery and Magnetic Hyperthermia in Cancer Therapy

**DOI:** 10.1101/2024.11.04.621819

**Authors:** Mohamadreza Tavakoli, Samane Maghsoudian, Amir Rezaei-Aderiani, Maliheh Hajiramezanali, Mahdiyar Amani, Elham Sharifikolouei, Mohammad Hossein Ghahremani, Mohammad Raoufi, Hamidreza Motasadizadeh, Rassoul Dinarvand

## Abstract

Due to the limited therapeutic efficacy and side effects associated with conventional chemotherapy, researchers have turned their attention to developing targeted drug delivery systems using advanced nanotechnology. Coating nanoparticles (NPs) with cell membranes is a promising strategy because it extends their circulation times and allows them to selectively adhere to damaged vessel sites through the platelet membrane surface, thereby enhancing tumor uptake. Herein, we have developed a biomimetic drug delivery system consisting of superparamagnetic iron oxide nanoparticles (SPIONs) coated by platelet membranes (PM) for carrying Paclitaxel (PTX) to exploit the synergism effect of chemotherapy and magnetic hyperthermia. Controlled-release PTX nanoparticles exhibited consistent behavior over time, indicating no significant difference in release between SPION/PTX and SPION/PTX/PM at pH 7.4. However, at pH 5.5, improved release was observed, specifically a 1.4-fold increase for SPION/PTX/PM. The confocal and flow cytometry results show an enhancement in the cellular uptake of SPION/PTX/PM nanoparticles, with an average fluorescence intensity of 142 ±12.5. MTT results indicated that SPION/PTX/PM demonstrates superior cytotoxic effects compared to SPION/PTX and PTX alone, showing an IC50 value of 5 μg/mL after 48 h of treatment. Furthermore, the IC50 decreased to 1 μg/mL when an alternating magnetic field was applied. Hence, the in vivo results and histopathological staining showed that the SPION/PTX/PM-AFM treatment group exhibited the highest rate of tumor growth inhibition, reaching nearly 92.14 %. These findings highlight the potential of using platelet membrane-coated nanoparticles for targeted delivery, combining magnetic hyperthermia and chemotherapy to minimize chemotherapy’s undesirable effects while maximizing therapeutic outcomes.

## 1. Introduction

Breast cancer is one of the most prevalent malignancies globally and the leading cause of cancer-related mortality among women. Current treatment options for breast cancer include surgical intervention, chemotherapy, and immunotherapy, used either alone or in combination(1, 2). Despite the critical role of paclitaxel in breast cancer treatment, several challenges persist, including toxicity, the necessity for high dosing, adverse side effects, and non-specific targeting. (3, 4). These issues underscore the need for ongoing research into more effective and targeted therapeutic strategies. The development of novel drug delivery systems and combination therapies may enhance disease management significantly.

Various carriers have been explored for the stable and targeted delivery of PTX, including micelles (2, 5), Albumin nanoparticles (NPs) (6), molecularly imprinted polymer NPs (7), and carbon nanotubes (8). Recent advancements in the application of superparamagnetic iron oxide nanoparticles (SPION) have shown promise across multiple fields, such as drug delivery (5, 9), gene delivery (10), magnetic resonance imaging (MRI) (11), and magnetic hyperthermia therapy (MHT) (12, 13). MHT is the sensitivity of tumor cells to increased temperatures. When SPIONs are exposed to a high-frequency magnetic field, magnetic hysteresis generates heat. This localized heating can raise the temperature of cancer cells to therapeutic levels, typically between 41 and 45 degrees Celsius, leading to various effects on these cells. These effects include alterations in the function of enzyme proteins, reduced membrane transport that impacts nucleic acid synthesis and DNA repair enzymes, and changes in DNA composition. Ultimately, this can result in cell death or increase the susceptibility of cancer cells to other treatments, such as chemotherapy and radiotherapy. The efficacy of SPION nanoparticles in MHT has been extensively investigated for treating several types of cancer, including prostate (14), breast(15) , and liver cancers (16).

Effective targeting and accumulation of SPIONs in tumors are crucial for the success of hyperthermia treatments (17). However, like many other nanomaterials, SPIONs face challenges related to their clearance from the bloodstream due to recognition by the reticuloendothelial system, even after being coated with biocompatible polyethylene glycol (18-20). This significantly hinders their accumulation and efficacy at the tumor site. Surface modification techniques, including tumor-targeting polymers (12) and antibody (21) or peptide sequences (22), aim to enhance tumor targeting efficiency and provide potential solutions to overcome this barrier. Nonetheless, opsonization and unintended clearance by the reticuloendothelial system remain significant challenges for these targeted systems (23).

Recently, the idea that natural cell membranes innately evade the immune system greatly inspired researchers to employ them in designing biomimetic drug delivery systems composed of cell membranes such as platelets, macrophages, white and red blood cells, and stem cells (24). Thanks to immune avoidance (25), prolonged blood circulation time (26), inflammatory site accumulation (27), and the capability to identify circulating tumor cells (28), platelet membranes (PM) have a great capability to be used as carriers of chemotherapeutics and NPs. The latest studies have revealed that PM can bind to specific tumor surface molecules and selectively target them (29-31). For instance, Li et al. demonstrated that mesoporous silica NPs coated with PM accumulated in tumor tissues via the targeted adhesion of PM surface to damaged vessels (32). In another study, Zhuang et al. designed a metal-organic framework nanoparticle covered with PM and showed substantial antitumor targeting (31). Hence, the advantageous features of PM make them an ideal candidate for incorporation into a drug delivery system specifically targeting tumors.

Despite the promising potential of SPIONs and the benefits of platelet membrane coatings, significant challenges remain unaddressed in optimizing the delivery of chemotherapeutics like PTX. there is limited understanding of their interactions with SPIONs and how these combined systems behave in dynamic biological environments (33). The synergistic effects of combining SPIONs’ magnetic targeting and hyperthermia capabilities with the immune-evasive properties of PM have not been fully investigated, particularly in the context of overcoming the high interstitial pressure and dysfunctional vasculature of tumors that limit nanoparticle penetration and retention (34, 35). Moreover, recent studies have highlighted the need for more in-depth evaluations of the pharmacokinetics, biodistribution, and long-term biocompatibility of these combined systems in vivo (34, 35). Addressing these gaps is critical for advancing the development of more effective and targeted chemotherapeutic delivery systems that can improve patient outcomes by minimizing side effects and enhancing drug efficacy at tumor sites.

The aim of this study is to develop and evaluate a novel drug delivery system that combines the chemotherapeutic efficacy of PTX with the targeting and hyperthermia capabilities of SPIONs and the immune evasion properties of PM. Specifically, the study focuses on the preparation, characterization, and in vitro and in vivo assessment of PTX-loaded SPIONs coated with PM (SPION/PTX/PM). By integrating these components, the study seeks to enhance the targeted delivery and therapeutic efficacy of PTX in breast cancer treatment, while minimizing systemic side effects and overcoming current challenges in nanoparticle drug delivery systems (schematic1).

**Schematic 1.**
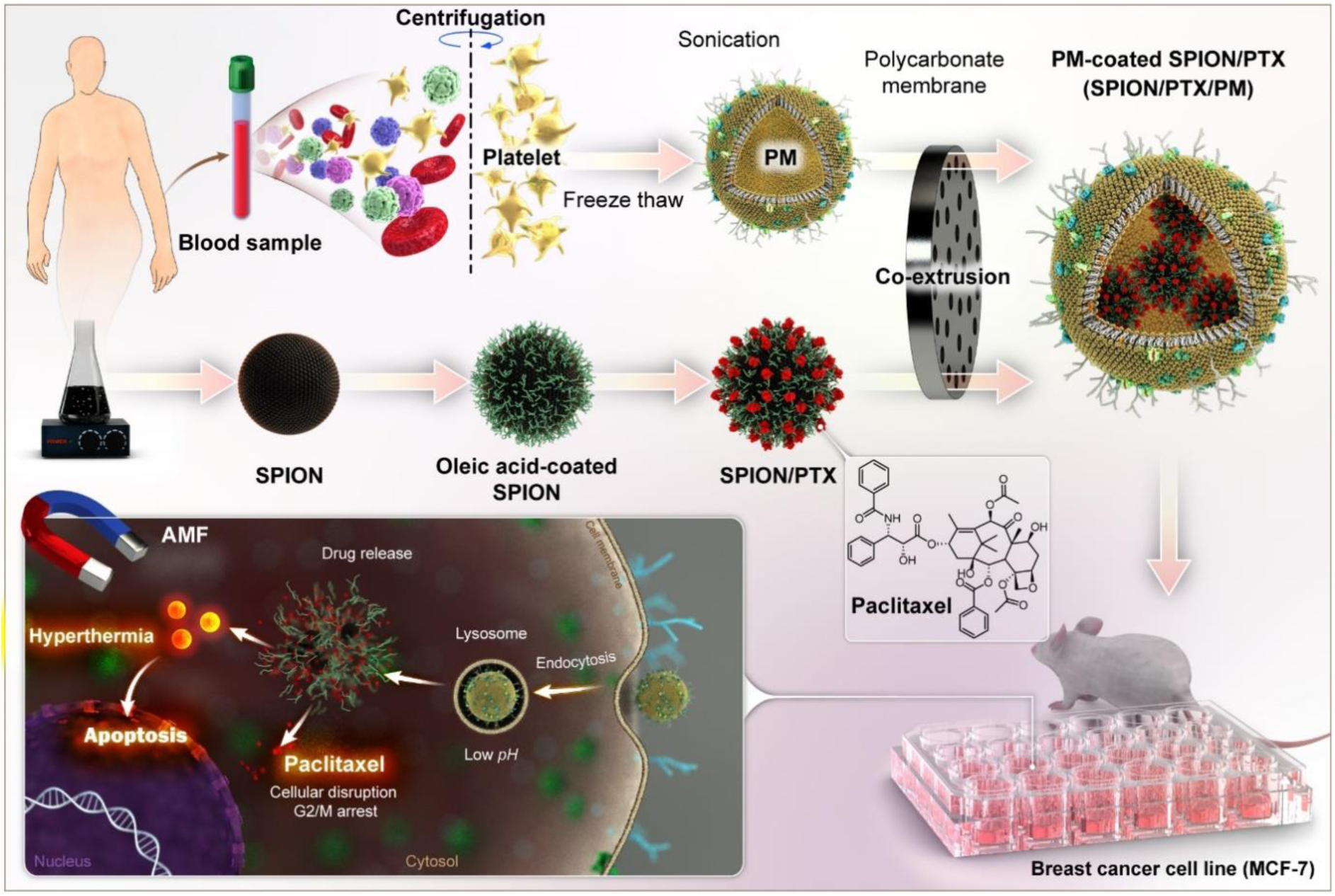
Schematic illustration of SPION/PTX/PM preparation from PM and PTX-loaded oleic acid-coated SPION by co-extrusion method. AMF: alternating magnetic field, G2: Gap 2 phase, M: Mitotic phase.

## 2. Materials and Methods

### 2.1 Materials

The ferrous chloride tetrahydrate (FeCl_2_.4H_2_O), ferric chloride hexahydrate (FeCl_3_.6H_2_O), Pluronic-F127, and Oleic acid (OA) were from Sigma Aldrich (USA). The acetonitrile (HPLC grade), Dimethyl Sulfoxide (DMSO), Ethylene diamine tetraacetic Acid (EDTA) Hydrochloric acid (HCL), Iron (III) nitrate nonahydrate (Fe(NO_3_)_3_⋅9H_2_O), HPLC grade Ethanol (99.5%), HPLC grade Acetone (99.8%) were from Merck (Germany). MCF-7 cell line was from Pasteur Institute, Tehran, Iran. 3-(4,5-dimethylthiazol-2-yl)-2,5-diphenyl tetrazolium bromide) (MTT), Cell culture media (DMEM) and fetal bovine serum (FBS) were from GIBCO (Life Technologies Inc., New York, USA). The 96-well microplate was from Greinerbio-one (Germany). Phenyl methyl sulfonyl fluoride (PMSF), Leupeptin, and Pepstatin were from Roche Diagnostics (Germany). The BCA protein assay kit was from Thermo Scientific (USA).

### 2.2 Preparation and characterization of SPIONs

SPIONs were synthesized by co-precipitation of Fe (III) and Fe (II) in a basic NH_4_OH aqueous solution (36). In brief, 3 mL of 5M NH_4_OH solution was added in dropwise to a mixture of 0.1 M for Fe (III) and Fe (II) under a nitrogen atmosphere for 1 min while the solution was constantly stirred. After adding 100 mg of OA, the solution was stirred for 20 min under nitrogen gas pressure. Afterward, while continuing stirring, the mixture was heated to 80℃ for 30 min. SPIONs coated with OA were separated by placing a strong magnet under the solution mixture for about 5 min. After discarding the supernatant, OA-coated NPs were washed thrice with deionized (DI) water. The nanoparticles were then dispersed in 45 mL of an aqueous solution containing 100 mg Pluronic F-127. Following an overnight stir on a magnetic stirrer, the magnetic NPs suspension was centrifuged (Sigma. 3-30K) for 15 min at 1,000 rpm and 10°C to eliminate the aggregates.

The average hydrodynamic size of NPs was determined by dynamic laser light scattering (DLS) in DI water (detection angle of 90°) at the wavelength of 633 nm (Nano-ZS; Malvern Instruments, Malvern, UK). The suspension prepared in DI water was used for measure zeta potential. The size and morphology of NPs were determined using transmission electron microscopy (TEM) (CEM 902A; Zeiss, Oberkochen, Germany). A diluted aqueous nanoparticle suspension drop was located onto a Formvar-coated 150 mesh copper TEM grid and then dried with air.

### 2.3 Preparation of PTX loaded SPION (SPION/PTX)

The method utilized for loading PTX into the OA-coating layer of the synthesized SPIONs was based on a previously established protocol (36). While stirring, different volumes from the ethyl alcohol solution of PTX (1 mg/mL) were added dropwise to an aqueous dispersion of OA-coated SPIONs (3.66 mg/mL). Stirring was continued overnight to complete the loading of PTX. The glass vial of the mixture was located next to a strong magnet for about 5 h to separate free PTX from SPION/PTX Next, SPION/PTX was washed twice by resuspending in DI water and then separated using a magnet, as mentioned earlier. To ensure the removal of free PTX, the concentration of PTX was assessed in each washing solution using the HPLC method. At last, SPION/PTX was dispersed in 5 mL of water.

### 2.4 Platelet membrane isolation

To efficiently obtain platelet membranes, fresh human platelet-rich plasma (PRP) (O^-^ blood type) in acid-citrate-dextrose was taken from the Iranian Blood Transfusion Organization. Before platelet collection, EDTA was added to a concentration of 5 mM. The platelets were separated from red and white blood cells by centrifugation at 100g for 20 min at room temperature. To ensure complete removal of the remaining blood cells, the resulting solution underwent another round of centrifugation, as mentioned. PRP was enriched with a PBS containing 1 mM EDTA and 2 μM prostaglandin E1 to inhibit platelet activation. Platelets were then separated by centrifugation at 800 rpm for 20 min at room temperature. After removing the supernatant, the obtained platelets were resuspended in PBS containing 1 mM of EDTA and protease inhibitors (PMSF 85 µg/mL, Leupeptin 0.5 µg/ml, and Pepstatin 0.7 µg/mL).

The PM was obtained from purified PRP through three freeze-thaw cycles. First, the purified PRP was frozen at -80°C and then thawed at room temperature for three consecutive cycles. The solution was centrifuged at 4000g for 3 min to pellet the PM, followed by washing with PBS (containing protease inhibitors). The isolated PM was then redispersed in DI water and sonicated for 5 min using a bath sonicator (Fisher Scientific FS30D) at 42 kHz and 100 W. After sonication, the PMs were extruded through 400 nm polycarbonate membranes ten times, and then through 200 nm polycarbonate membranes to produce platelet vesicles. DLS and TEM were used to verify the formation of platelet vesicles.

### 2.5 BCA assay

The protein concentration in the solution of platelet vesicles was determined using the Pierce® BCA protein assay kit (37). Initially, the standard bovine serum albumin (BSA) solution was prepared as per the kit instructions. Then reagents A and B were then mixed based on the protocol. Subsequently, both the standard and unknown samples were transferred into a 96-well microplate (n=3), followed by the addition of working reagents to each well. The 96-well microplate was then placed on a shaker for 30 s to ensure thorough mixing and then incubated for 30 min. Lastly, using a microplate reader (BioTek ELx800), the absorbance of each well was measured at 562 nm.

### 2.6 Preparation of SPION/PTX/PM

The extrusion method was used to coat PM around prepared SPION/PTX. The SPION/PTX and PM were combined at a weight ratio of 2:1 and sequentially co-extruded ten times through 400 nm and 200 nm polycarbonate membranes to obtain the NPs encapsulated by PM. Finally, the resulting NPs were isolated using a magnet.

### 2.7 Drug loading (DL) and entrapment efficiency (EE) of SPION/PTX and SPION/PTX/PM NPs

The amount of PTX loaded onto SPION/PTX was assayed using the indirect method. After separating the NPs from free PTX with a magnet, the solution surrounding the NPs was isolated, freeze-dried, and then dissolved in acetonitrile. The solution was centrifuged at 14000 g for 10 min to eliminate any remaining NPs. The supernatant was then assayed for PTX using the HPLC method (pump (Wellchrom, K-1001, Knauer, Berlin, Germany), UV detector (Wellchrom, K-2600, Knaur), Column: C18 (4.6×250 mm i.d., pore size 5 µm, Knauer, Berlin, Germany), mobile phase: Acetonitrile and DI water (55:45)). Finally, the amount of loaded PTX was calculated by subtracting the initially added drug amount from the obtained amount. The percentage of loaded drug and its encapsulation efficiency was calculated according to formulas (1) and (2).

To determine the amount of PTX loaded onto SPION/PTX and SPION/PTX/PM, direct method was employed. After separating the nanoparticles from free PTX using a magnet, the nanoparticles were dissolved in acetonitrile. The solution was then centrifuged to remove the nanoparticles, and the supernatant was analyzed by HPLC (pump (Wellchrom, K-1001, Knauer, Berlin, Germany), UV detector (Wellchrom, K-2600, Knaur), Column: C18 (4.6×250 mm i.d., pore size 5 µm, Knauer, Berlin, Germany, mobile phase: Acetonitrile and DI water (55:45). The drug loading percentage and entrapment efficiency were calculated using formulas (1) and (2).

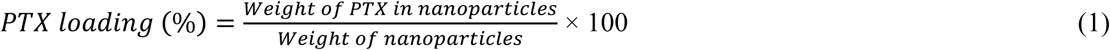

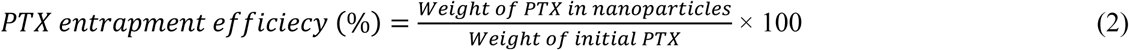

### 2.8 characterization of SPIONs

Magnetic properties of SPIONs and SPION/PTX/PM were determined using a vibrating sample magnetometer (Model 155; Princeton Applied Research, Oak Ridge, TN, USA) at room temperature under a magnetic field up to 10 kOe. Fourier transform infrared (FTIR) spectra were obtained using a Nicolet 6700 spectrometer (Tensor27; Bruker; Germany) to determine the chemical structure of all synthesized NPs. X-ray diffraction (XRD) measurements were conducted on SPIONs using a Siemens D5000 X-ray diffractometer (Siemens, Germany) with Cu Kα radiation (λ = 1.5406 Å). The scan range of 2Ɵ was from 0 to 80°. The results were analyzed by Diffract-plus software.

### 2.9 In vitro PTX release

*In vitro* PTX release from SPIONs and SPION/PTX/PM was determined using the dialysis method. Briefly, a 1-mL suspension of SPIONs and SPION/PTX/PM was placed in a dialysis bag with a 6-kD cutoff, then immersed in 10 mL of PBS containing 1% (w/v) Tween-80 as the release medium and incubated at 37°C with stirring at 100 rpm. The drug release profile was evaluated in PBS at two different pH levels (7.4 and 5.5). Samples of 1 mL were replaced with fresh medium at specified intervals, and the concentration of PTX in the supernatant was measured using HPLC after centrifugation at 17,000 x g for 10 min.

To assess The release profile of PTX under the influence of an alternating magnetic field (AMF), different formulations were placed in 6 kDa dialysis bags immersed in release media at pH 7.4. While the formulations were mildly stirred at 25 °C under an AMF (60 Gs) for 1 h, the samples were taken from the medium at predetermined time intervals (5, 10, 15, 20, 30, 45, and 60 min).

### 2.10 In vitro cytotoxicity study

The MTT assay was used to evaluate the *in vitro* cytotoxicity of free PTX, SPION/PTX, SPION/PM, and SPION/PTX/PM in MCF-7 cells. The MCF-7 breast cancer cell line was cultured in DMEM supplemented with 10% FBS and 1% penicillin-streptomycin at 37°C with 5% CO_2_. The cells were seeded in a 96-well plate at a density of 5 × 10^3^ cells per well. After 24 h incubation, medium was replaced with DMDM containing free PTX, SPION/PTX, SPION/PM, and SPION/PTX/PM. following incubating for 24 and 48 h, 50 µL of MTT solution (1 mg/mL) was added to each well and further incubated for another 4 h. Formazan were dissolved with DMSO, the absorbance was measured at 540 nm by a microplate reader (BioTek ELx800).

A separate cell plate was prepared using the same method mentioned above to evaluate the toxicity of SPION/PTX, SPION/PM, and SPION/PTX/PM in the presence of AMF. The cells were exposed to NPs, after 4 h subjected to AMF (19.5 kA/m and 389 kHz), and then incubated for 24 h.

### 2.11 Cellular uptake study

The cellular uptake of SPION, and SPION/PM, was investigated using confocal laser scanning microscopy (CLSM) and flow cytometry. initially, Fluorescein isothiocyanate (FITC) was loaded onto SPIONs and SPION/PM NPs following the method described earlier for PTX loading in dark ambient. cells were seeded in 6-well plates at 2.0 × 10^5^ cells per well and incubated for 24 h. Subsequently the original medium was replaced with media containing FITC, SPION/FITC, and SPION/FITC/PM (with FITC concentration of 250 µg/mL). after 4 h of incubation, cells were washed with PBS and detached using Trypsin. The cell suspension was centrifuged at 1000 rpm for 5 min to collect cells which were then resuspension in 200 μL of PBS and analyzed using a flow cytometer to measure the average fluorescence intensity.

For CLSM analysis, MCF-7 cells were cultured on 24-well plate microscope slides at 5 × 10^4^ cells per well for 24 h. Next, after adding SPION/FITC or SPION/FITC/PM to the cells, followed by a 2-h treatment period. After removing the culture medium, cells were washed thrice with ice-cold PBS. The cells were then fixed with 4% paraformaldehyde solution for 5 min, and the cell nuclei were stained with HOECHST 33258 (10 µg/mL) for 20 min. After three PBS washes, fluorescent images of the cells were captured via an LSM710 CLSM (Nikon Inc., USA).

### 2.12 In vivo tumor inhibition study

For in vivo studies on the anti-cancer efficacy of nanoparticles, 1.5 × 10^6^ 4T1 cells suspended in 200 µL of PBS were injected subcutaneously into the flanks of female Balb/c mice. Once the tumors reached volumes between 50-100 mm^3^, the mice were randomly assigned to one of five groups: SPION/PTX/PM-AFM, SPION/PTX/PM, PTX, SPION/PM, and a control group (3 mice per group). Subsequently, samples were intravenously injected into the mice four times on days 0, 3, 6, and 9. Over a 18-day period, daily monitoring included assessing changes in body weight and tumor volumes. In the AFM-receiving group, mice were exposed to AFM (19.5 kA/m and 389 kHz) for 1 h, starting 4 h after each injection.

Tumor volume was calculated using the formula: “Tumor volume = (w^2^ × l) / 2” where ’l’ and ’w’ denote the length and width of the tumor, respectively.

### 2.13 Histopathological analysis

After 21 days of starting the treatment, the mice were euthanized. Tumor tissues and major organs were fixed in 4% formalin for 24 h and then transferred to paraffin for preservation. Thin sections (3-5 µm) of the tissues were prepared on slides and stained using the Hematoxylin and Eosin (H&E) protocol to achieve effective nuclear and cytoplasmic staining. The semi-quantitative analysis was conducted by a pathologist using parameters such as mitotic index (MI), nuclear pleomorphism (NP), and apoptosis.

### 2.14 Statistical analysis

Origin software was used to compare the results obtained from experiments, graphs, and other statistical methods. The test results were presented as mean ± SD. Statistical analysis results with a P value of less than 0.05 were considered significant.

## 3. Results and Discussion

### 3.1 Characterization of the NPs

In the first step, X-ray diffraction was conducted on the synthesized SPIONs to confirm their presence. The diffraction pattern aligns with the crystal structure of Fe₃O₄, indicating successful synthesis. As shown in Figure 1a, characteristic peaks corresponding to Fe₃O₄ are observed at diffraction angles of 2θ = 30.2°, 35.5°, 43.0°, 53.5°, 57.1°, and 62.5°, which correspond to the lattice planes (220), (311), (400), (422), (511), and (440), respectively.

**Figure 1.**
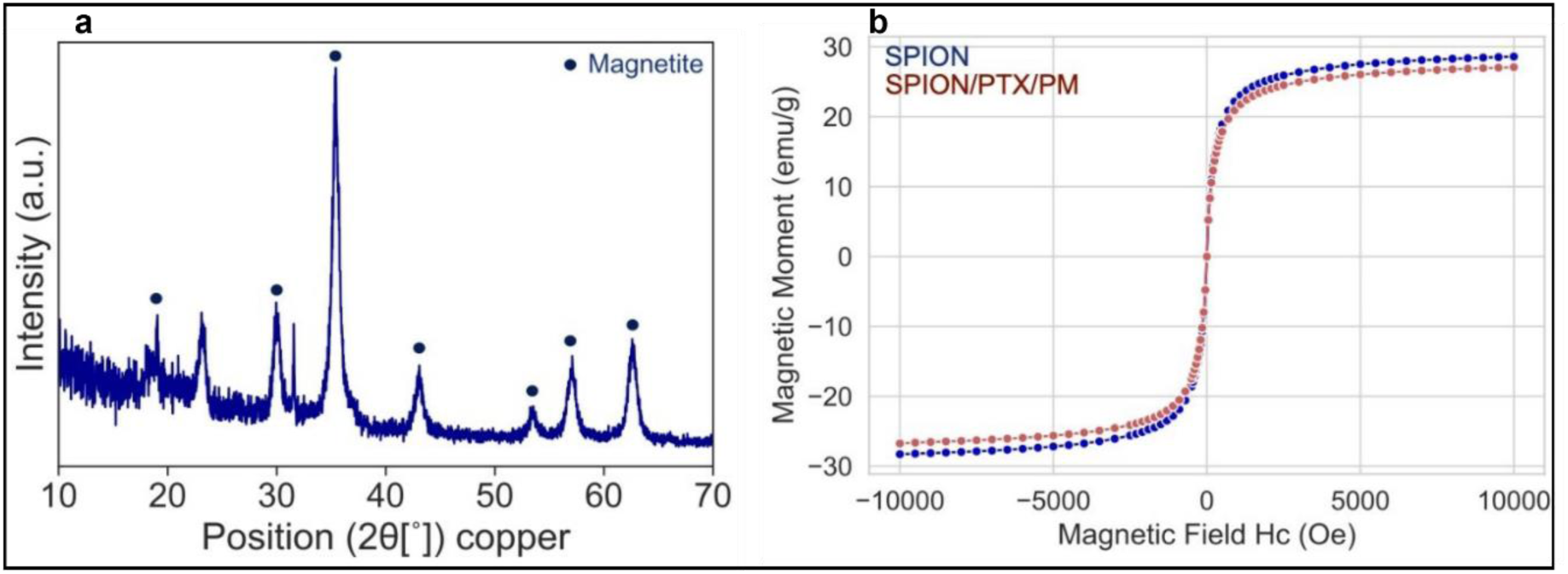
a) XRD analysis of synthesized SPIONs. Magnetite (Fe3O4) was detected as the primary crystalline phase; b) Magnetization analysis of SPIONs and SPION/PTX/PM conducted by VSM.

After modifications such as membrane coating, SPIONs must retain their magnetic properties, essential for hyperthermia induction and MRI. Therefore, magnetization vs applied magnetic field were performed on both coated and uncoated SPIONs using a vibrating sample magnetometer (Fig. 2b). The results demonstrate that the superparamagnetic character of SPIONs remains unchanged after PM coating.

**Figure 2.**
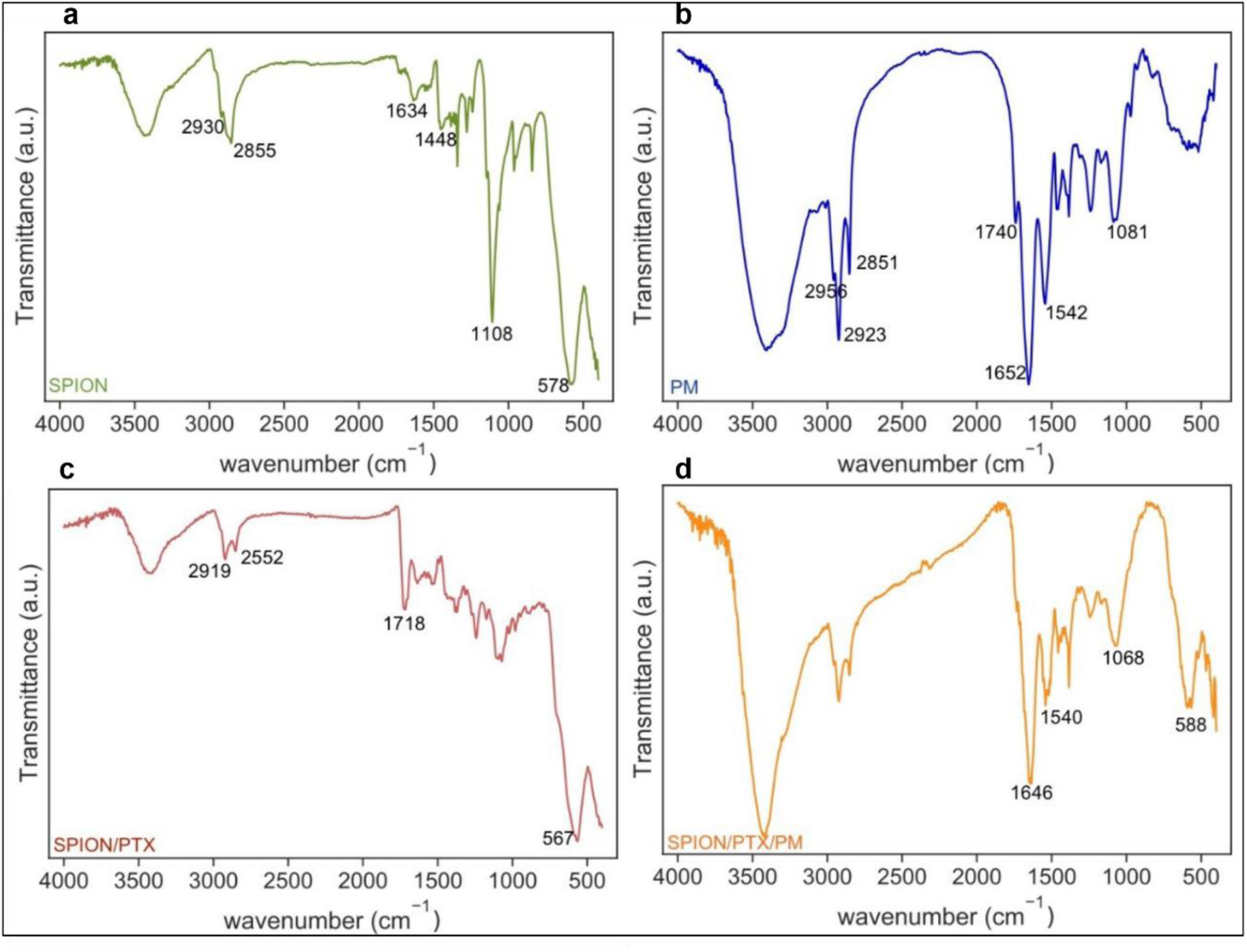
FTIR of a) SPIONs, b) PMs, c) SPION/PTX, and d) SPION/PTX/PM.

FTIR was performed to confirm the synthesis of SPION/PTX/PM. In this regard, the FTIR spectra was obtained for uncoated SPIONs, PM, SPION/PTX, and SPION/PTX/PM; and are presented in Figure 2. The FTIR spectrum of synthesized SPIONs (Fig. 2a) shows an adsorption peak in the region of 578 cm^-1,^ indicating the vibration of the Fe-O bond and the formation of the crystal structure of Fe_3_O_4_ NPs. In addition, the band in the range of 3600-3000 cm^-1^ corresponds to O-H bond vibrations. The presence of oleic acid on the surface of NPs was confirmed by CH_3_ vibrations observed at 2855 cm^-1^ and 2930 cm^-1^. Previous studies indicate that the C=O vibration of the carboxyl group typically appears at 1708 cm^-1^ in the spectrum of pure liquid oleic acid, which was absent in the synthesized SPIONs. Instead two new bands at 1448 cm^-1^ and 1634 cm^-1^ emerged, characterized by symmetric and asymmetric COO^−^ stretching vibrations. The strong peak at 1108 cm^-1^ also indicates the stretching of a single C-O band. Therefore, the presence of carboxylate and CH_3_ vibration confirmed the presence of oleic acid in the synthesized sample, chemically adsorbed through its carboxylate group onto the surface of SPION.

In the FTIR spectrum of PMs (Fig. 3b), the band at 1740 cm^-1^ corresponds to the carbonyl group of the ester structure found in the lipids comprising PM. Peaks at 2956, 2923, and 2851 cm^-1^ represent asymmetric and symmetric vibration of CH_2_ and CH_3_ groups within the bilayer structure of PM, respectively. The Vibration observed at 1652 cm^-1^ shows the C=O carbonyl bond in the amide group of protein, and 1542 cm^-1^ shows the vibration of the N-H bond and the C-N in the structure of the proteins. The symmetric vibration of P=O bond of PO_2−_ in the phospholipids structure of PM appears at 1081 cm^-1^ (38) .

**Figure 3.**
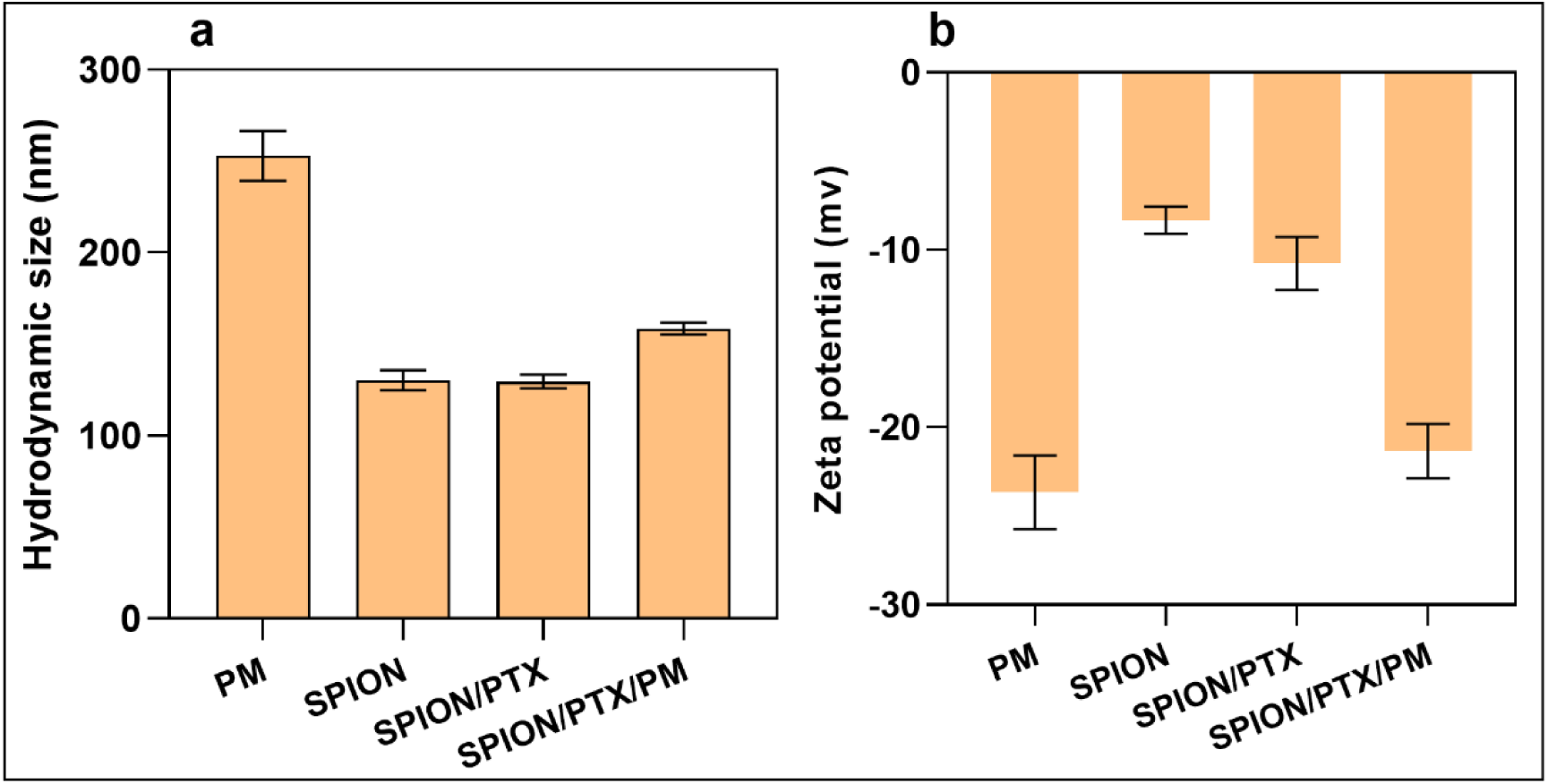
a) Size distribution of PM; SPIONs; SPION/PTX; SPION/PTX/PM and b) zeta potential of PM; SPIONs; SPION/PTX; SPION/PTX/PM by DLS.

**Figure 4.**
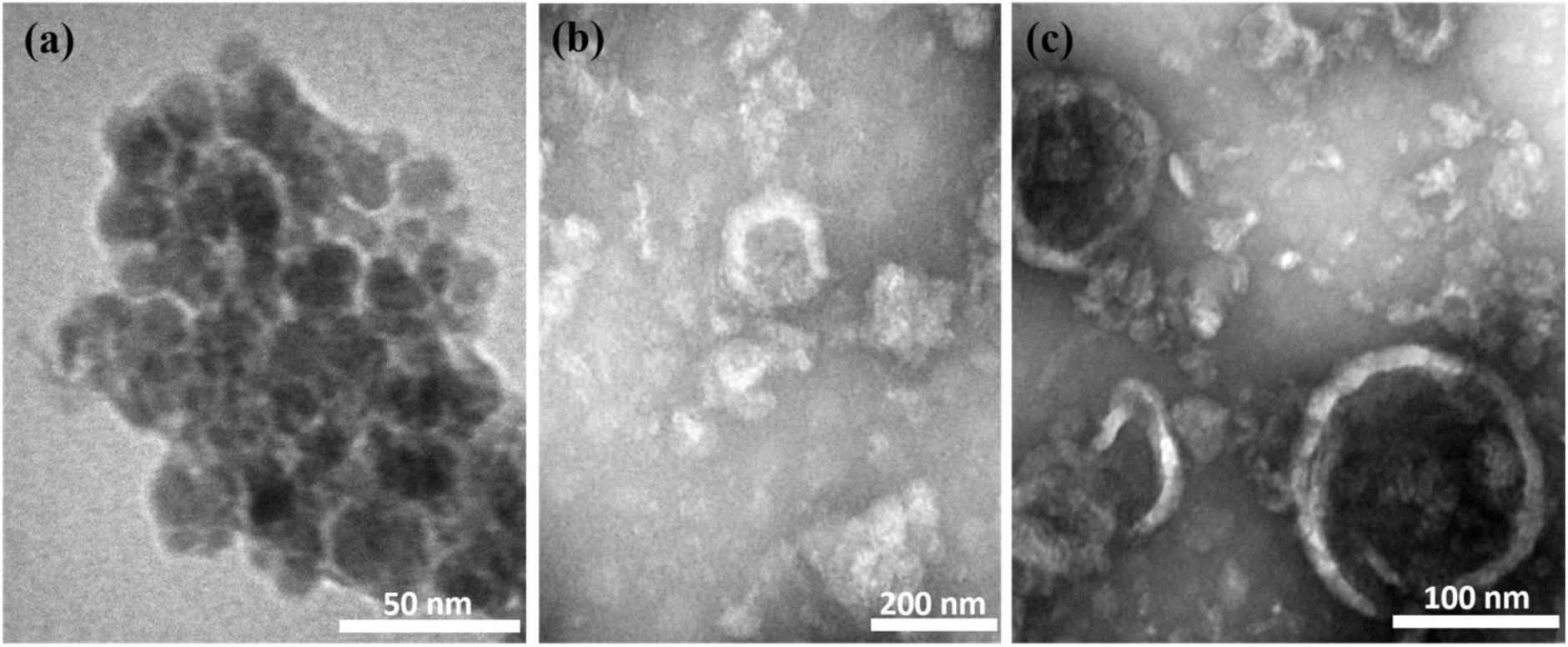
TEM bright field images of a) SPIONs, b) PM, and c) SPION/PTX/PM

In the FTIR spectrum of SPION/PTX (Fig. 2c), the peak at the 567 cm^-1^ confirms to the Fe-O band characteristic of magnetite, confirming the presence of SPIONs in SPION/PTX. The vibration at 1718 cm-1 corresponds to the C=O ester bond present in PTX, , indicating successful loading of PTX in within SPIONs structure (39).

The FTIR spectrum of SPION/PTX/PM NPs (Fig. 2d) shows a peak at 588 cm^-1^ identical to SPIONs. alongside new absorption peaks at 1068 cm^-1^ related to the P=O band vibration, 1540 cm^-1^ related to C-N band vibration and 1646 cm^-1^ related to C=O amide band vibrationin proteins. These peaks were absent in SPION/PTX, indicating the presence of PM coating SPION/PTX. Upon internalization of SPION/PTX within PM, the characteristic peaks of PTX diminish, such as the strong absorption peak at 1718 cm^-1^. Moreover, the FTIR spectra of SPION/PTX/PM resemble those of platelet cell membranes, suggesting cell membranes are present on the surface of SPION/PTX/PM. Thus, as illustrated by FTIR results, the surface of SPION/PTX is efficiently covered by PM.

Our results align with Kazarian et al.’s approach (40) using infrared spectroscopy to study biological systems. Their study revealed distinctive bands associated with organic compounds (CH) and common peptides (N-H, O-H, C=O, C-O) present in samples with platelet coatings. The bands were notably enhanced in the platelet-containing samples, indicating the presence of PM.

The next step involves evaluating the amount of platelets coated onto the SPIONs. The concentration of platelet proteins in the SPION/PM sample was determined using the BSA kit, yielding a result of 600 µg/mL. This result indicated a coating efficiency of 79.45%. The measurement was based on absorbance values obtained from BSA standards at known concentrations. Particle size, PDI, and zeta potential were measured using DLS. As shown in Fig. 3a, PM has a diameter of 226 nm and a zeta potential of -24 mV (PDI 0.185), consistent with previous reports on platelet vesicle sizes and zeta potentials(31, 41). SPIONs have a hydrodynamic diameter of 125 nm and a zeta potential of -9 mV (PDI 0.441). SPION/PTX has a hydrodynamic diameter of approximately 128 nm and a zeta potential of -12 mV (PDI 0.082). Finally, DLS measurements of SPION/PTX/PM reveal an increased average size of 147 nm and a more negative zeta potential of -22 mV (PDI 0.170), indicating effective coating with platelets. The surface charge of SPION/PTX/PM (−22 mV) is similar to that of PMs (−24 mV), suggesting that the surface properties of SPION/PTX/PM are comparable to those of the platelet membrane.

To visually confirm these findings, TEM was used to examine the morphology of SPION, PM, and SPION/PTX/PM (Fig. 5). Fig. 5b shows spherical PM vesicles characterized by a bilayer cell membrane, visible as a white halo. Notably, TEM images of SPION/PM reveal distinct membrane coatings on the SPIONs with white halo. (Fig. 5c). According to Hu et al., the white halo observed around spherical nanoparticles indicates the presence of a cellular membrane layer on these particles (42).

**Figure 5.**
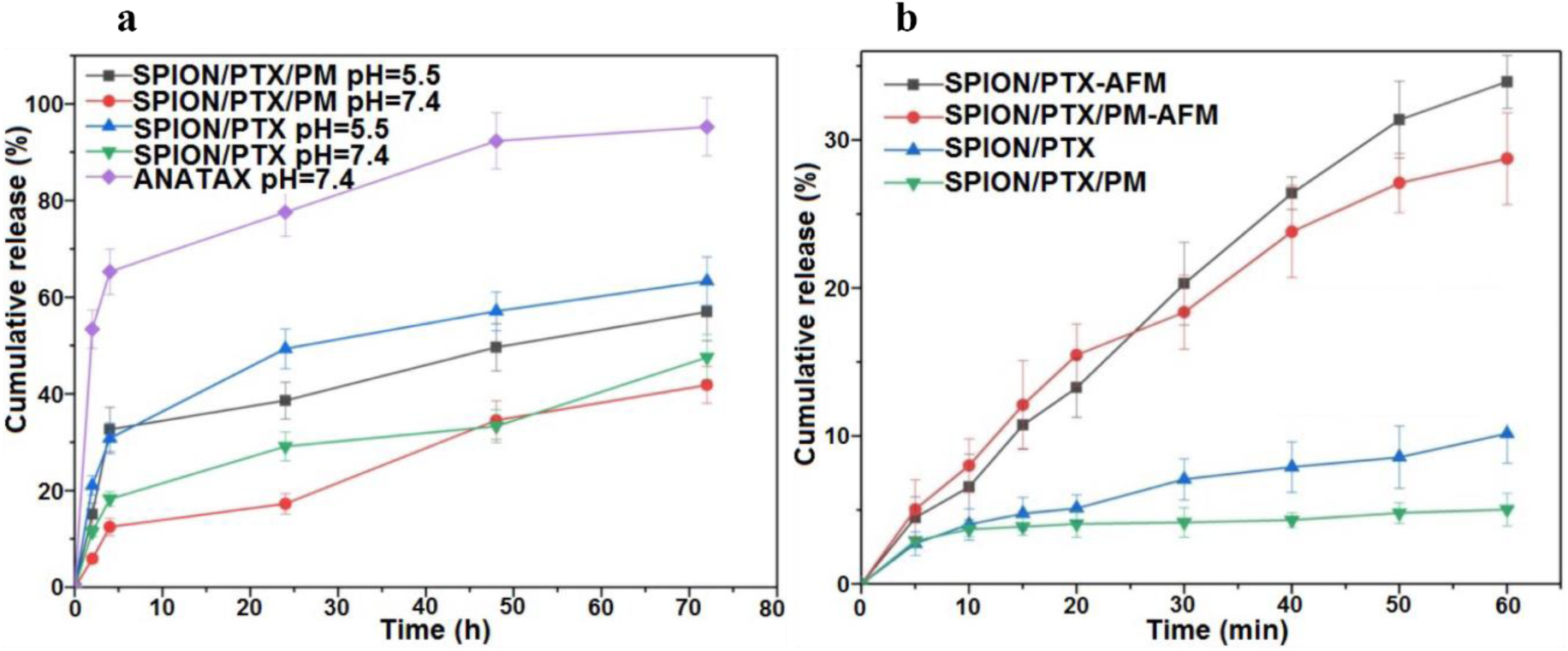
a) PTX release from SPION/PTX and SPION/PTX/PM in pH=7.4 and Ph= 5.5 compared with commercial PTX (Anzatax), **b)** PTX release from SPION/PTX and SPION/PTX/PM in pH=7.4 in the presence and absence of AMF.

### 3.2 Drug loading and encapsulation efficiency

To examine the impact of different weight ratios of PTX to NPs on DL content and EE in SPION/PTX NPs, different volumes of a PTX solution (10 mg/mL) and an SPION suspension (3.66 mg/mL) were combined. The results for EE and DL are presented in Table 1. NPs synthesized with a weight ratio of 3:11 showed the highest EE (90.01%), whit a DL of 19.7%. Therefore, this ratio was selected for further experiments.

**Table 1.** The effect of different weight ratios of PTX to SPIONs on DL and EE of SPION/ PTX

| weight ratio of PTX<br>to SPIONs (w/w) | EE (%) | DL (%) |
| --- | --- | --- |
| 0.55 : 11 | 60.83 | 2.95 |
| 1.50 : 11 | 69.33 | 8.63 |
| 2.20 : 11 | 73.31 | 12.78 |
| 2.50 : 11 | 76.93 | 14.88 |
| 3.00 : 11 | 90.01 | 19.71 |
| 3.30 : 11 | 84.70 | 20.28 |

To analyze the DL and EE of SPION/PTX/PM NPs, the unloaded PTX did not separate from SPION/PTX prior PM coating, aiming to enhance EE through PTX’s strong protein binding and the presence of membrane proteins. After isolating free drug from the final SPION/PTX/PM NPs using a magnet, the EE and DL of resulting NPs were measured at 95.5% and 14.5%, respectively.

### 3.3 Drug release study

The cumulative release of various formulations at pH 5.5 and 7.4 was assessed, with the commercial product Anzatax used as the control. As shown in Fig. 6, both groups demonstrated an initial burst release of PTX within the first 4 h. Under pH = 5.5, 32% and 30% of PTX were released from SPION/PTX and SPION/PTX/PM, respectively, while under pH=7.4, the releases were 12% and 18%. Over the following 68 h, PTX release continued at a slower rate. As illustrated in Fig. 5a, after 72 h, only 41% and 47% of PTX were released in pH=7.4, whereas 57% and 63% were released under pH= 5.5 from SPION/PTX and SPION/PTX/PM, respectively. These results indicate a stable and prolonged release of PTX from these nanoparticles. Notably, SPION/PTX/PM exhibited better drug release control compared to SPION/PTX. Specifically, PTX release in the SPION/PTX/PM group was 1.4 times higher in pH = 5.5 than pH= 7.4. Acidic conditions enhance the efficiency of cancer drug delivery due to the lower pH of tumor cells, which helps minimize damage to normal cells. (42). Similar PTX release behavior has been observed in other nanocarriers, such as lipid-coated NPs (43), stem cell membrane-coated PLGA NPs (44), and gelatin nanogels (45), primarily due to the hydrophobic properties of PTX.

**Figure 6.**
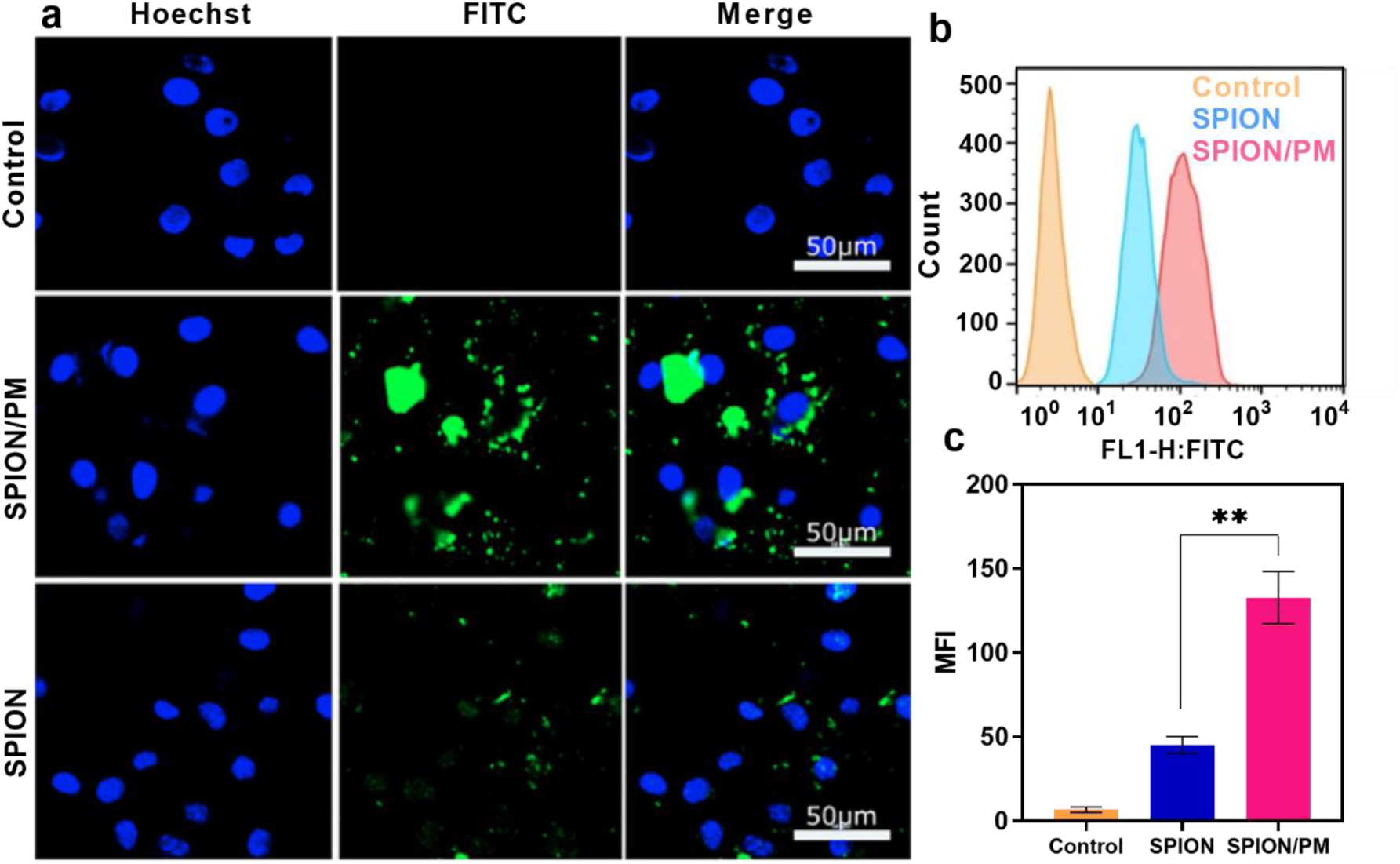
In vitro assessment of SPION/FITC and SPION/FITC/PM uptake into MCF-7 cells using CLSM the blue color shows the nuclei of cells stained with Hoechst, and green corresponds to NPs containing FITC ((Scale bar: 50 μm, Magnification 40×)). (b) Quantitative MCF-7 cellular uptake of SPIONs compared with PM-SPIONs and control group (PBS). NPs are loaded with FITC. (c) Mean florescent intensity (MFI) of different of samples against MCF-7 cells.

The cumulative release of PTX from SPION/PTX under exposure to an AMF was assessed using a similar method. As shown in Fig. 6b, the release of PTX from SPION/PTX and SPION/PTX/PM increased by 3.3 and 5.7 times, respectively, after 60 min with AMF compared to without it. This increase is due to the breakdown of hydrophobic bonds between PTX and OA layers surrounding the SPIONs, as well as the disruption of the plasma membrane structure caused by the heat generated from SPIONs under AMF. AMF converts magnetic energy into thermal energy in superparamagnetic particles (46-48).

### 3.4 Cellular uptake studies

CLSM was performed to evaluate the effect of PM coating on the uptake of NPs by MCF-7 cells, as depicted in Fig. 6. NPs were loaded with FITC to enable visualization of cellular uptake. the CLSM image of MCF-7 cells showed that green fluorescence from SPION/FITC/PM exhibited higher intensity than that of SPION/FITC. This reveals an enhancement uptake of the SPIONs by MCF-7 cells due to the presence of PM coating.

Furthermore, flow cytometry was utilized to quantitatively evaluate the cellular uptake of SPION /FITC/PM. The flow cytometry graph (Fig.6b, c) reveals that MCF-7 cells incubated with PM-coated SPION/FITC NPs exhibited a mean fluorescence intensity (MFI) of 142 ± 12.5, whereas those treated with uncoated SPION/FITC showed a MFI of 46 ± 9.8. Membrane coating substantially augmented the cellular uptake of SPIONs, as the fluorescence intensity of SPION/FITC/PM was about 3.1-fold of SPION/FITC after 4 h of incubation. Therefore, it can be concluded that the presence of PM on SPIONs facilitates their entry into MCF-7 cancer cells, probably due to the targeting ability conferred by surface proteins of the PM, which facilitate their interaction with tumor cells (30).

### 3.5 In vitro cytotoxicity

The MTT assay was conducted to evaluate the cytotoxicity of SPION/PTX, SPION/PTX/PM, SPION/PM, and PTX against MCF-7 cells (With and without AFM), with results shown in Fig. 8. Up to a concentration of 5 µg/mL, all samples and free PTX maintained over 50% cell viability after 24 hours. This cell viability percentage in PTX-containing nanoparticles may be due to a hydrophobic interaction between PTX and the OA shell, leading to gradual PTX release from the nanoparticles, as shown in the release results (Fig. 5a). Fig. 8a indicates a significant difference (p<0.05) in cell death rates between SPION/PTX/PM with PTX and SPION/PTX only at a concentration of 10 µg/ml, likely due to increased cellular uptake of SPION/PTX/PM.

**Figure 7.**
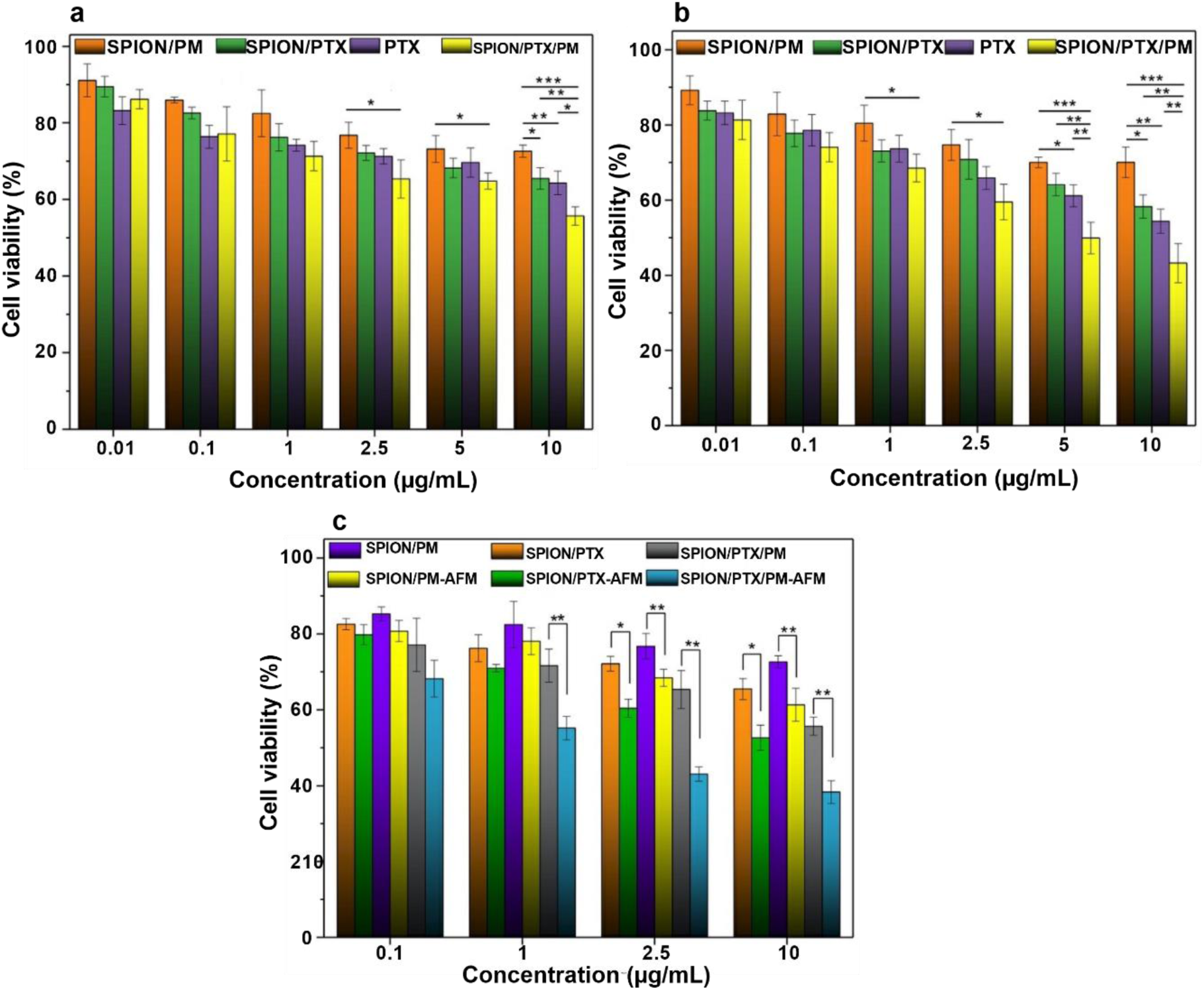
MTT results of free PTX, SPION/PTX, SPION/PM, and SPION/PTX/PM at various concentrations of PTX on MCF-7 cells after **a)** 24 h , **b)** 48 h and **c)** after 24 h with and without AMF

**Figure 8.**
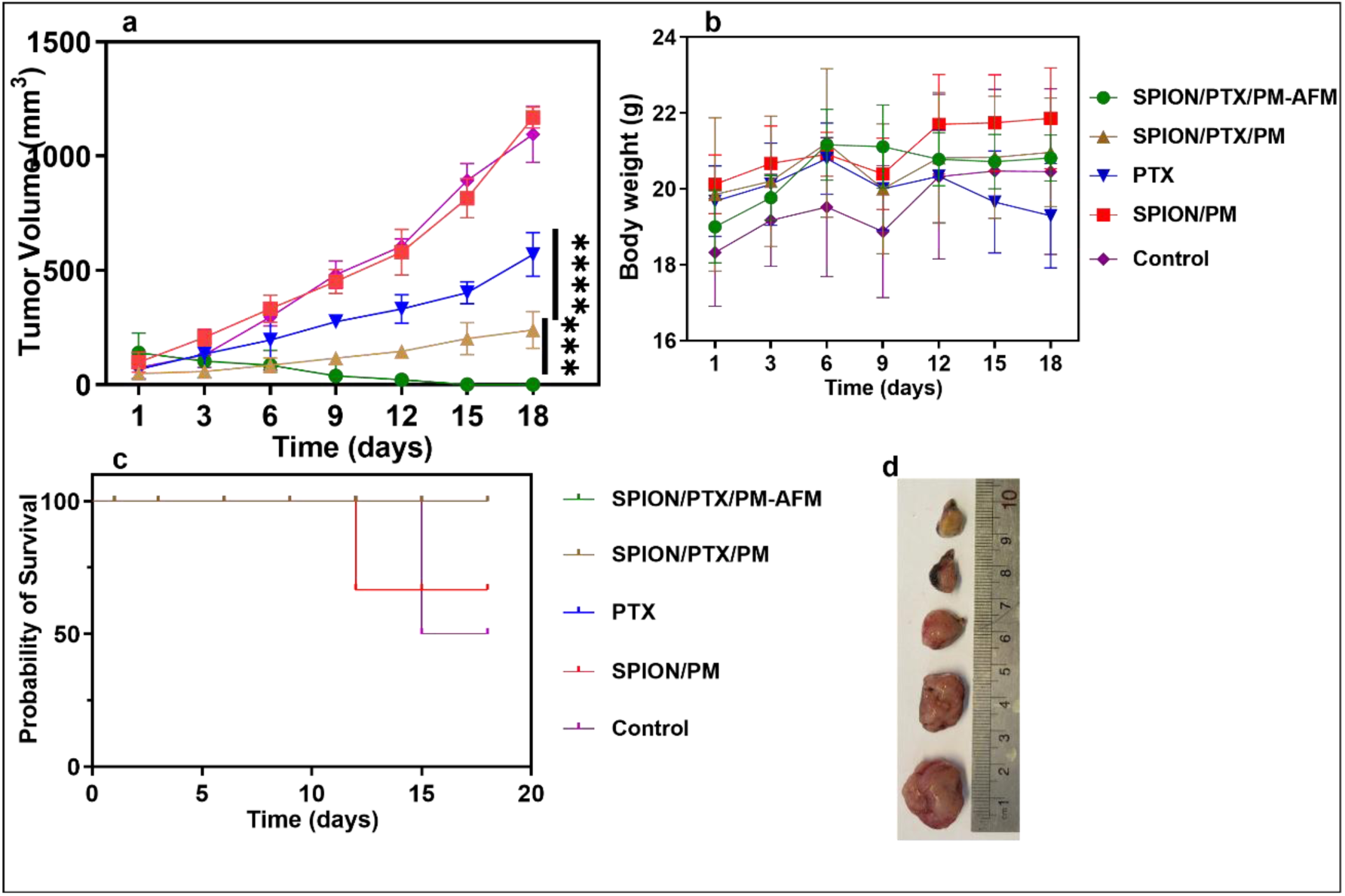
In vivo performance of different formulation in mice bearing 4T1 tumors, (a) Tumor volume chart of treated groups with saline, SPION/PTX/PM, SPION/PTX/PM-AMF, PTX, and SPION/PM (****p < 0.0001 and (***p < 0.001) (b) fluctuation of body weight in all groups, and (c) Survival rates of tumor-bearing mice in different group. (d) The size of tumors removed from mice after 18 days.

After 48 h (Fig. 7b), SPION/PTX/PM exhibited the highest cell death, indicating greater absorption compared to other nanoparticles and free PTX. The estimated IC50 for SPION/PTX/PM nanoparticles at 48 h was approximately 5.21 µg/mL. Cell mortality in other groups did not reach 50% at any concentration within the 48-h period. SPION/PM nanoparticles showed 15 % cell death even at the highest concentrations, indicating their low toxicity. At concentrations of 5 and 10 µg/mL, there was a significant difference (p<0.05) in cell death rates between SPION/PTX/PM and SPION/PTX, likely due to increased cellular uptake with PM, consistent with CLSM flow cytometry findings.

After 24 h of incubation with SPION/PTX/PM without AMF, maximum cytotoxicity was observed, reaching 51.78 % at a concentration of 10 µg/mL. However, exposure to AMF resulted in an IC50 of 5 µg/mL, indicating significant therapeutic magnetic effects by AMF (Fig. 7c). In both coated and uncoated nanoparticles, the cell death rate increased in the presence AFM compared to its absence. This effect can be attributed to several phenomena related to heat production by superparamagnetic nanoparticles in the presence of AMF, as cancer cells are highly sensitive to temperature increases (49). In the presence of AMF, SPION-containing NPs generate heat. When the temperature exceeds 40°C, cancer cells are susceptible to damage to the membrane and cytoskeletal integrity, potentially initiating apoptosis (50). Additionally, high temperatures damage the fluidity of the cell membrane, which may enhance the uptake of SPION/PTX/PM nanoparticles into the cells (51). Furthermore, the heat generated by SPIONs in the presence of AMF could disrupt the hydrophobic bond between PTX and the OA shell, leading to increased drug release (46).

Ultimately, SPION/PTX/PM caused greater cytotoxicity in the presence of AMF at all concentrations compared to SPION/PTX/AFM, likely due to increased cellular uptake of surface proteins in the cell membrane (52).

In summary, the cytotoxic effect of SPION/PTX/PM in the presence and absence of AMF was superior to unirradiated SPION/PTX/PM and unirradiated SPION/PM, respectively. Thus, combinatorial chemotherapy and hyperthermia therapy in SPION/PTX/PM NPs considerably restrained MCF-7 cell proliferation.

### 3.6 In vivo study

The study evaluated the anti-tumor effect of NPs in 4 tumor-bearing mice over an 18-day period post-initial injection. Mice received intravenous treatments of NP/P/PTX, NP/P, saline and free PTX on days 0, 3, 6, 9 Throughout the study, mice in different experimental groups were monitored daily for signs of distress, tumor volume changes, and body weight fluctuations. As depicted in Figure 8a, mice treated with SPION/PTX/PM showed a significant reduction in tumor volume 18 days after the initial injection compared to those treated with free PTX. By day 11, two mice in the SPION/PTX/PM group had achieved complete remission, with the remaining mice in this group demonstrating significantly lower tumor volumes compared to the free PTX group (****p < 0.0001). No deaths occurred during the study period, indicating an enhanced anti-tumor activity of NP/P/PTX NPs compared to free PTX. Moreover, no clinical signs of PTX-related side effects such as nutrition or mobility issues were observed during the treatment. However, it is notable that significant weight loss was observed only in the PTX-treated group from day 12 onwards (Figure. 8b). The groups treated with free PTX exhibited a notable increase in tumor volume from day 9 onwards, reaching 2.41 times the initial tumor volume by the end of the study, which was significantly different from the SPION/PTX/PM group. The mortality rate in these groups was one death per day by day 12 (Figure. 8c). Mice treated with SPION/PTX/PM/AFM displayed slower tumor progression through day 18, with two mice achieving complete remission by day 11. From day 12 onwards, tumor volume in this group significantly diverged from the SPION/PTX/PM group (***p < 0.001). No significant weight loss was observed until the mice were euthanized on day 18 and their tumors were examined. The control group and SPION/PM recipient group exhibited the fastest tumor progression and highest tumor volume at the end of the study. The SPION/PM group had a mortality rate of one death per day by day 12, while the control group experienced one death by day 15.

These findings clearly emphasize the effective impact of SPION/PTX/PM and suggest that this innovative approach holds promise for more efficient cancer therapies by reducing systemic toxicity. The research shows that coating NPs with platelets enhances anti-tumor activity through improved targeting, immune evasion, prolonged circulation, increased penetration, and better retention (53, 54). Moreover, there are reports indicating that platelets interact with circulating tumor cells (CTCs) through various surface receptors and signaling molecules. When NPs are enveloped with PM, they can adopt these interactions (55). This implies that SPION/PM can directly bind to tumor cells and deliver their payload more effectively, potentially hindering the spread of CTCs (metastasis).

### 3.7 H&E analysis

To evaluate whether there was a positive anti-tumor effect in the treatment groups compared to the PBS control group, various parameters (mitotic count (MI), nuclear pleomorphic (NP), and apoptosis) were analyzed using H&E-stained images of tumors from three mice per group, reviewed by a pathologist (Figure. 9a). Ten photographs were captured from each tumor sample using a 40x objective lens (10 HPF). Effective treatment resulted in increased apoptosis and reduced mitosis in pathological assessments. The results showed that the SPION/PTX/PM-treated group had the lowest mitotic count compared to the other groups (Figure. 9d). Increased nuclear pleomorphism score, characterized by enlarged nuclei indicating cells primed for division, was most pronounced in the PBS group and significantly lower in the SPION/PTX/PM-AFM-treated group (Figure. 9b). Enhanced therapeutic effects were indicated by increased apoptotic cells following PTX uptake by tumor cells and increased connective tissue. Apoptosis and therapeutic effects were notably higher in the SPION/PTX/PM-AMF and SPION/PTX/PM-treated group compared to the control (Figure. 9c), highlighting extensive areas of connective tissue that support the anticancer efficacy of these NPs (56).

**Figure 9.**
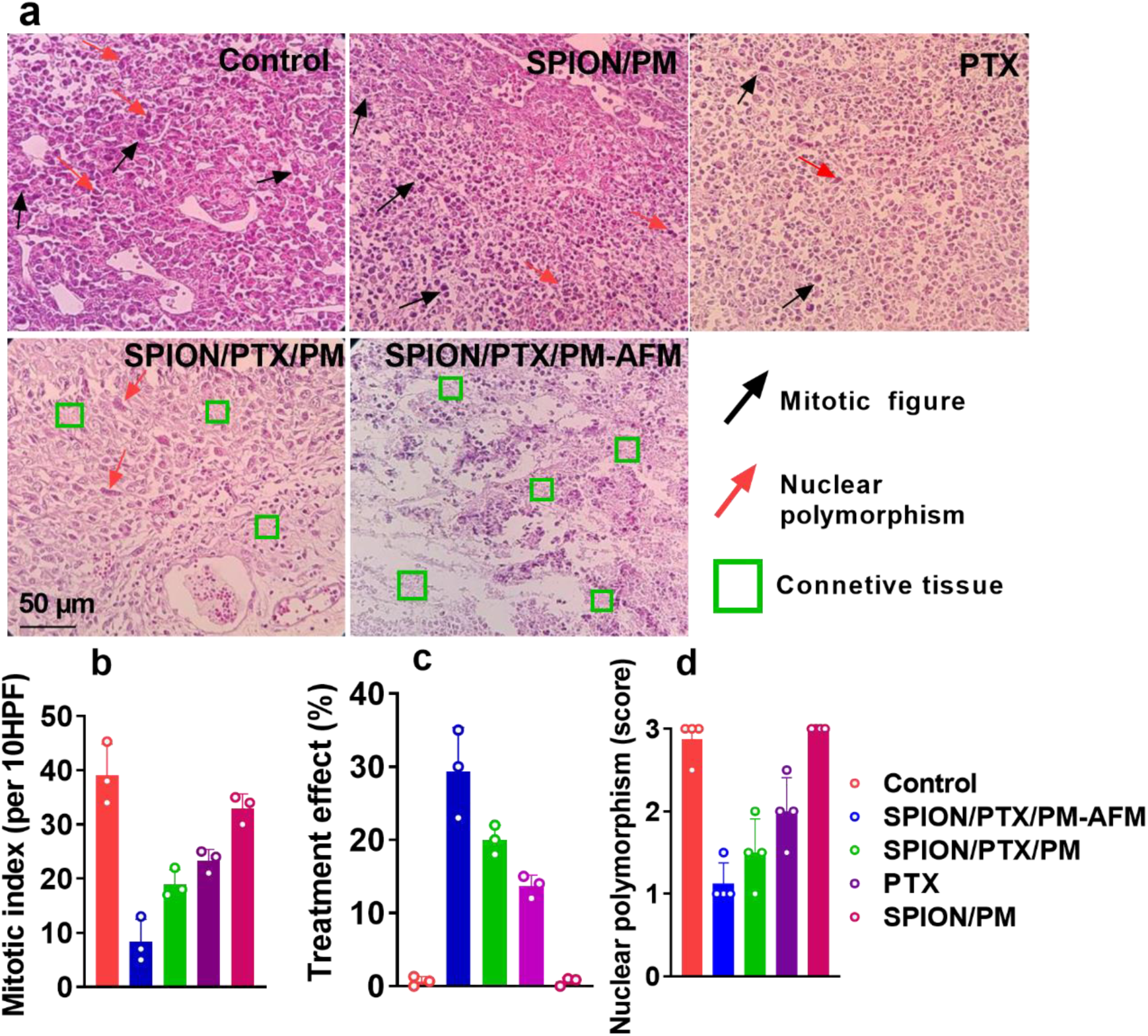
(a) H&E staining of breast tumor in all groups after 18 days (Magnification 40×. Scale bar: 50 μm), semiquantitative analysis of H&E staining tumoral slices (b) Treatment effect, (c) MI, and (d) NP. Data are presented as mean ± SD (n =3).

## 4. Conclusions

By combining chemotherapy and magnetic hyperthermia, we successfully engineered a nano-sized biomimetic drug delivery platform to suppress the growth of MCF-7 cells. The study suggests that our system can prevent the sudden release of PTX, making it an ideal choice for drug delivery systems targeting tumor tissues. Also, due to the superior uptake of SPION/PTX/PM by MCF-7 tumor cells compared to SPION/PTX without PM coating, using our system in vivo may reduce the side effects of PTX. The *in vitro* cytotoxicity assay indicated that the MCF-7 cells incubated with SPION/PTX/PM in the presence of AMF obtained the IC_50_ of 1 μg/mLand had a significantly lower growth rate than those without AMF. In conclusion, with the combination of magnetotherapy, chemotherapy, and PM targeting ability, SPION/PTX/PM can be considered a hopeful biomimetic drug delivery for future breast cancer therapy.

## Acknowledgment

This research was a part of Mohammadreza Tavakoli’s Pharm.D thesis, supported by a grant from the Nanotechnology Research Centre, Tehran University of Medical Sciences.

## Ethics

All experiments were performed in accordance with the ethical guidelines of Tehran University of Medical Sciences.

